# A fully automated spike sorting algorithm using t-distributed neighbor embedding and density based clustering

**DOI:** 10.1101/418913

**Authors:** Mohammad Hossein Nadian, Saeed Karimimehr, Jafar Doostmohammadi, Ali Ghazizadeh, Reza Lashgari

## Abstract

In this study, a new spike sorting method was developed based on a combination of two methods, t-Distributed Stochastic Neighbor Embedding (t-SNE) and Density-Based Spatial Clustering of Applications with Noise (DBSCAN). Parameters of both methods were simultaneously optimized using a Genetic Algorithm (GA) using a simulated dataset containing 2 to 20 simultaneously recorded neurons. The performance of this method was evaluated using both the simulated dataset as well as real multichannel electrophysiological data. The results indicated that our fully automated algorithm using t-SNE-DBSCAN outperforms other state-of-the-art algorithms and human experts in spike sorting especially when there are a large number of simultaneously recorded units. Our algorithm also determines the noise waveforms and has an overall high sensitivity, precision and accuracy for correctly classifying waveforms belonging to each neuron (all >90%) without the need for manual corrections afterwards. Our method can be a crucial part of the analysis pipeline in particular when manual sorting of units is becoming prohibitive due to the sheer number of recorded neurons per session.

## 1. Introduction

Neurons are the building blocks of the nervous system. Inspecting and investigating the activity of single neurons is the foundation for understanding brain mechanisms. For a few decades, it has been possible to decode the behavior of a single neuron’s activity from multiunit brain recordings by measuring reflected current flows in the extracellular medium (Moser and Moser, 2013; O’Keefe, 1976; Olshausen and Field, 1997). Usually, neural action potentials or spikes are detected through extracellular recordings, typically using micro-electrodes (metal, silicon or glass micropipettes)(Buzsáki et al., 2012). The procedure of discriminating the spikes of each neuron from the multiunit recorded neural signal is usually referred to as “spike sorting” and often uses the shape of the spikes as discriminating information (Gibson et al., 2012). The goal in spike sorting is to determine the number of neurons recorded in a single channel and to specify the activation timing of each neuron (Quiroga, 2012).

Spike sorting has been studied extensively over the last few decades, with applications in neural interfaces (Oliynyk et al., 2012; Todorova et al., 2014), new prosthetic control devices (Zaghloul and Bayoumi, 2015), neurorehabilitation (Farina et al., 2013), and studies of cognitive function (Moser and Moser, 2013; O’Keefe, 1976). Review papers by Lewicki (1998), Gibson et al. (2012) and Rey et al. (2015) summarize a large number of techniques used for spike sorting. Some algorithms are designed for sorting single channel extracellular signals, while others were developed for recording systems like stereotrodes or tetrodes. Commonly applied methods for single-channel spike sorting are principal component analysis (PCA) (Adamos et al., 2008), Bayesian approaches (Haga et al., 2013), wavelet-based algorithms (Kim and Kim, 2003; Quiroga et al., 2004), and filter-based methods (Calabrese and Paninski, 2011). Spike sorting methods for multi-channel recordings have been proposed by Carlson et al. (2014); Swindale and Spacek (2014); and Rossant et al. (2016). Recently, electrode technology has made it possible to record from hundreds of neurons concurrently on a sub-millisecond timescale (Stevenson and Kording, 2011). However, the algorithm developments for spike sorting on this level have been much slower, making the efficient and reliable sorting of a large number of neurons challenging (Pedreira et al., 2012; Rey et al., 2015). Unfortunately, the amount of information that needs to be processed is now too high for spike sorting in a manual or semi-automated fashion (Einevoll et al., 2012). Thus, the main challenge is in developing automatic spike sorting algorithms (Wood et al., 2004).

In this paper, we introduce a novel automatic spike sorting algorithm based on combination of t-Distributed Stochastic Neighbor Embedding (t-SNE), developed by Maaten and Hinton (2008), and Density-based spatial clustering of applications with noise (DBSCAN), specifically designed for decoding a large number of neurons in single-channel extracellular recordings. The t-SNE algorithm, like other dimensionality reduction methods, transforms the data from high-dimensional space to a space of fewer dimensions. However, unlike most of the techniques, t-SNE is capable of retaining both the local and the global structures of the data in a single map (Maaten and Hinton, 2008).

## 2. Materials and methods

In this study, spike sorting performance with increasing number of neurons was evaluated using simulated 10 minute-long extracellular recording datasets including different numbers of single units, from 2 to 20 recorded on a single channel (total of 95 channels) (http://bioweb.me/CPGJNM2012-dataset). The spike sorting algorithm was also tested on real multi-electrode array neural recordings (Ghazizadeh et al., 2012). The neural recordings were conducted in Long–Evans rats from the Nucleus accumbens shell, using a drivable 16 electrode array. This system allowed recording simultaneously from multiple neurons across the 16 channels of recording (Ghazizadeh et al., 2012).

The overall procedure of the proposed spike sorting algorithm is illustrated in Fig 1. The method consists of the following steps:

1. Spike detection
2. Dimensionality reduction of spikes by t-SNE method
3. Clustering spikes by DBSCAN method

**Fig. 1.**
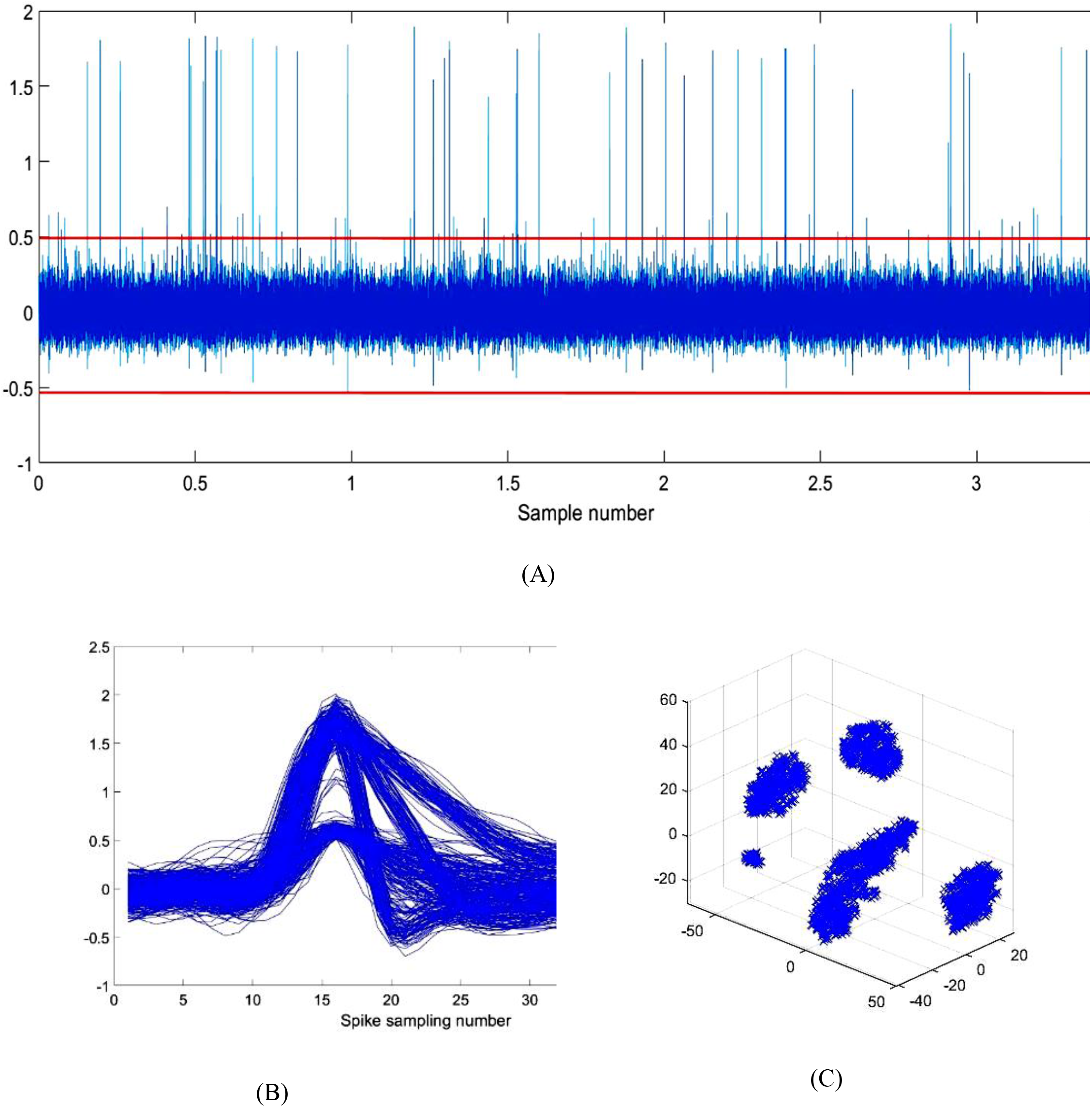
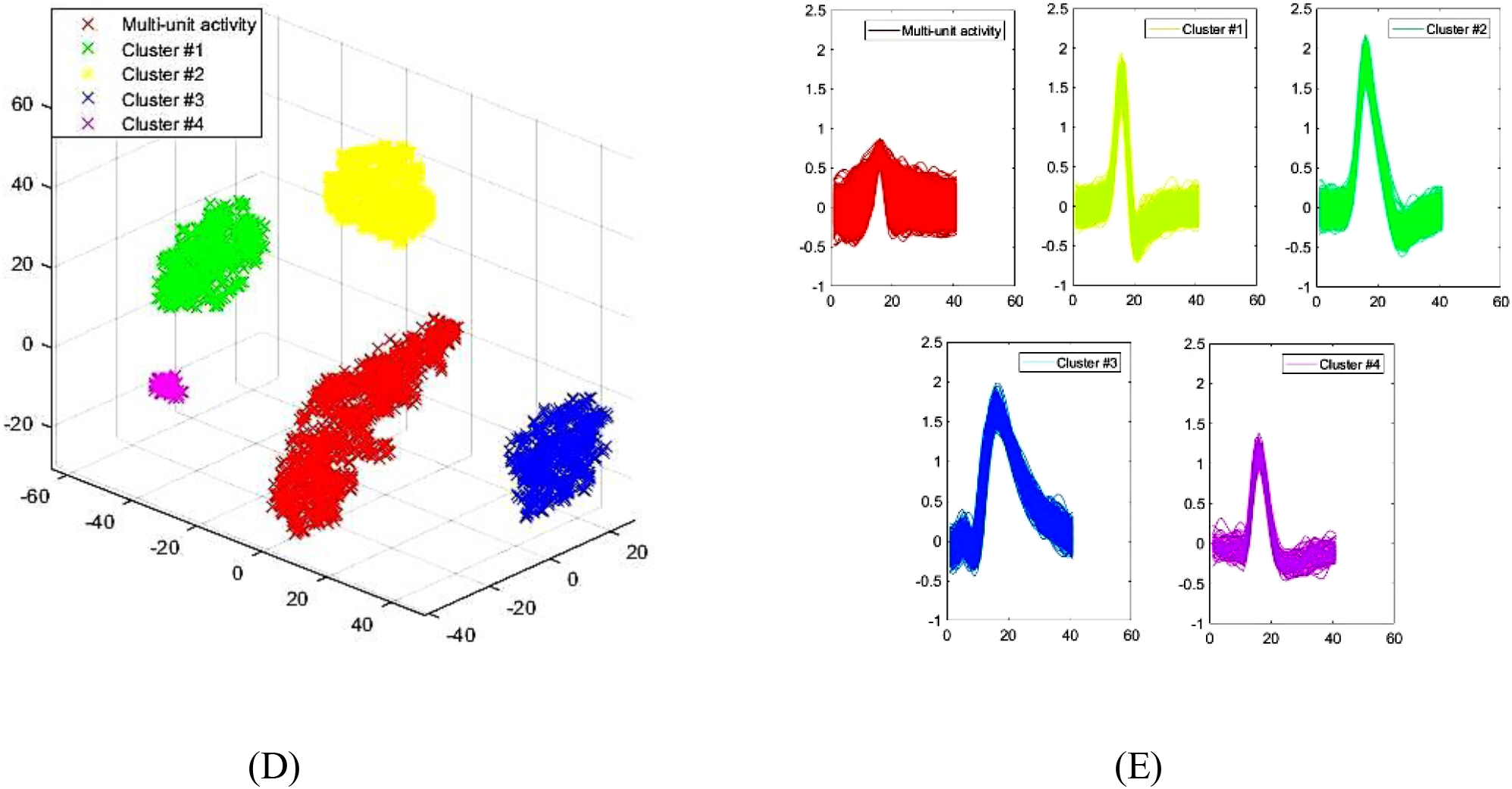
Shematic of consecutive steps for our automatic spike sorting procedure. (A) spikes are detected using an amplitude threshold after filtering raw data. (B) all of the detected spikes are extracted and aligned by their positive peaks. (C) dimensionality reduction was used in order to reduce complexity of clustering using t_SNE. (D) clustering algorithm using DBSCAN. (E) spike shapes associated with each cluster shown in D.

### 2.1. Spike detection

As all spike sorting algorithms, the initial step prior to the sorting method is to extract the spikes from the recording data. Primary preprocessing and band-pass filtering (300–6000 Hz, four pole Butterworth), enhances the spike detection on top of the background noise activity. Generally, spike detection is carried out by amplitude thresholding (T). To set an automatic threshold, a method is described based on the median absolute deviation (MAD).

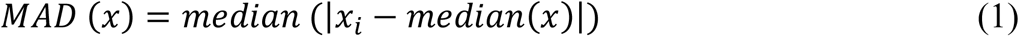

where *x* is the bandpass-filtered signal. In common cases where the median of signal (x) is zero, the Eq. 1 simplifies to:

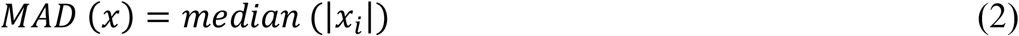

This method measures the variability of a univariate sample of quantitative data. Therefore, the variance is then robustly estimated as (Donoho and Johnstone, 1994):

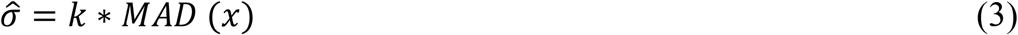

where k=1.4826 is a scale factor for normally distributed data. Generally, the amplitude threshold (T) is defined as the multiple (≅ 4) of an estimate of the standard deviation of the noise (Quiroga et al., 2004):

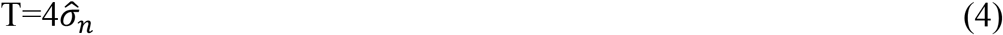

where *σ*̂_n_ is an estimate of the standard deviation of the background noise.

After detection of all of the likely spikes, the next step is to store the detected spike waveforms (∼2 ms long) in an array and align them to the spike peak such that the peaks are located in the middle of the array. This array is passed to the next step.

### 2.2. t-Distributed Stochastic Neighbor Embedding (t-SNE)

Before the clustering step, dimensions of spikes were reduced using the t-SNE method. Briefly, the t-SNE method converts high-dimensional data into a lower dimension by minimizing the Kullback-Leibler (KL) distance between the joint probability distribution defined between each two data points under high- and low-dimensional spaces. The joint probability distribution is made using a Gaussian centered at each point in the high dimensional space and a heavy-tailed Student-t distribution in the low dimensional space. A major strength of t-SNE is its capability in retaining the local structure of the data while also revealing some important global structure (Maaten and Hinton, 2008). The primary results of the t-SNE spikes map indicate that much of the local structure of the spikes is captured as well. Therefore, this method was used as a feature extraction algorithm for spikes. In order to achieve the best results, the parameters of the t-SNE algorithms must be optimized. One of the important parameters in the t-SNE algorithm is the distance metric which can be Euclidean, Standardized Euclidean, City block, Chebychev, Minkowski, Mahalanobis, Cosine, Linear Correlation, Spearman’s rank correlation, Hamming or Jaccard coefficient distances for both Gaussians and t-distributions. Also, perplexity or effective number of local neighbors of each point is another important parameter of the Kullback-Leibler algorithm which must be specified. Other parameters include exaggeration which is the size of natural clusters in the data, and number of dimension of the representation. All of the parameters were optimized using the Genetic Algorithm (GA) for simulated data (see below for details of GA).

#### 2.3. DBSCAN Clustering

Density-based spatial clustering of applications with noise (DBSCAN) is a density-based clustering method first introduced by Ester et al. (1996). In contrast to clustering methods like k-means, DBSCAN does not require one to specify the number of clusters in the data. Further, DBSCAN can find general cluster shapes and does not force all points to fall into detected clusters like the k-means algorithm. DBSCAN has two free parameters: size of neighborhood considered around each point (ε) and minimum number of points that should be in a cluster (MinPts). These parameters were also optimized along with the t-SNE parameters using GA for the above-mentioned ground truth dataset.

#### 2.4. Genetic Algorithms (GA)

The six free parameters of the algorithm (distance metric, perplexity, exaggeration, number of dimension, ε and MinPts) were optimized using a GA. Some options of the GA are indicated in Table 1. Six-dimensional strings (chromosome) are presented for solving this problem. The overall procedure for combined optimization of t-SNE and DBSCAN using GA, illustrated in Fig 2, includes the following steps:

**Table 1.** Some options for the GA solver.

| Parameters: | Value |
| --- | --- |
| Population type | Double vector |
| Population size | 200 |
| Creation function | Constraint function |
| Selection function | Stochastic uniform |
| Mutation function | Gaussian |
| Crossover function | Constraint dependent |
| Crossover fraction | 0.7 |
| Elite count | 5 |
| Generations | 100 |
| Function tolerance | $10^{-6}$ |
| Nonlinear constraint tolerance | $10^{-6}$ |

**Fig. 2.**
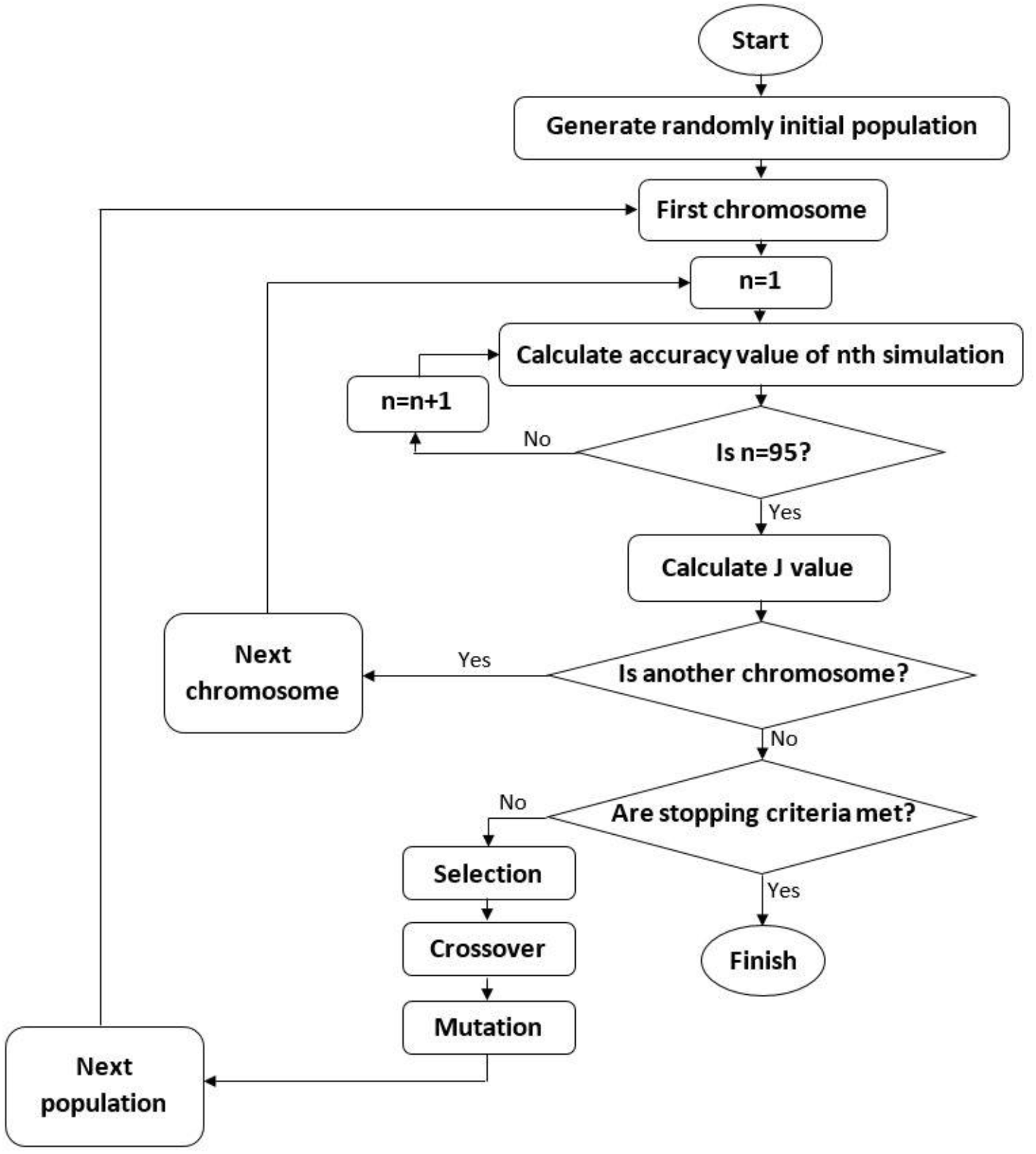
The flowchart of GA optimization procedure: 1) An initial population is generated by randomly choosing parameters from a specified range. 2) The accuracy of confusion matrix (Acc) for the simulated dataset is calculated. 3) The desirability function (J) was calculated based on the averages accuracy values of confusion matrixes across the sessions. 4) The process is repeated for all chromosomes in the pool. 5) Population is assessed and a new gene pool is formed by: recombination and mutation. 6) This procedure continues until either of stopping criteria.

1. The first step in the functioning of a GA is the generation of an initial population;If the initial population to the GA is good, then the algorithm has a better possibility of finding a good solution. The initial generation is random in the specified range and seemed to end up with acceptable results in pilot tests (Table 2)

**Table 2.** The initial population values of GA algorithm.
2. The accuracy of confusion matrix (Acc) is calculated for each session of simulated spikes that ranged from 2 to 20 neurons in a given session for every 6-dimensional string (chromosome) using Equation 5:

| Parameters: | limit |  |
| --- | --- | --- |
|  | Low | High |
| t-SNE: |  |  |
| Distance Metric* | 0 | 11 |
| Perplexity** | 30 | 150 |
| Exaggeration* | 3 | 6 |
| Number of Dimension* | 2 | 3 |
| DBSCAN: |  |  |
| $\epsilon$ *** | 1 | 10 |
| MinPts *** | 1 | 10 |
| * Step by increments of 1 (integer) |  |  |
| ** Step by increments of 10 (integer) |  |  |
| *** (floating point) |  |  |

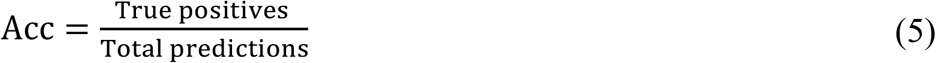
3. A cost function (J) was defined based on the average accuracy values (A) of confusion matrices of all sessions:

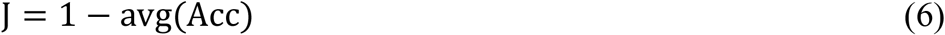
4. Scores were generated for each member of the current population by computing their fitness value. (population size = 200). This step selects parents based on their expectation.
5. Children are produced either by making a random vector from a Gaussian distribution to the parent (mutation) or by creating the child as a random weighted average of the parents (crossover). The crossover fraction of 0.7 means that 70% of children other than elite individuals are crossover children. Finally, a new population is replaced to form the next generation.
6. This procedure continues until either of the stopping criteria (Generations=100; Function tolerance=10^-6^; Nonlinear constraint tolerance=10^-6^) is reached.

#### 2.5. Evaluation of the optimal t-SNE and DBSCAN algorithm

In order to describe the performance of the designed classification model, a confusion matrix of each evaluated signal was obtained based on comparing the dataset ground truth with clusters identified by the t-SNE and DBSCAN algorithms. Each identified cluster was assessed as valid if at least 50% of its spikes were time-locked to the spikes of an actual simulated neuron (Martinez et al., 2009). The values of true positives (TP), true negatives (TN), false positives (FP) and false negatives (FN) of each confusion matrix were then calculated. Then, the values of sensitivity or true positive rate (TPR) and precision or positive predictive value (PPV) were obtained by the following equations:

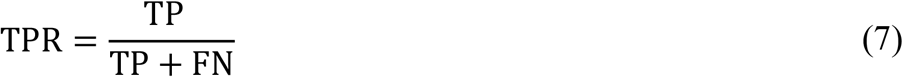

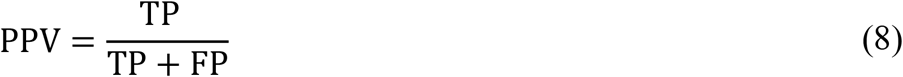

Another performance measure is based on the histogram of the Inter-Spike-Interval (ISI). When spikes from two different neurons are incorrectly classified as a single cluster, it is possible to have spikes with ISIs below the minimum refractory period of the neuron (taken to be below 2 ms in this study). If the proportion of refractory period violations is significant, it can be seen as a measure of poor isolation of the single units. Finally, the optimal t-SNE and DBSCAN spike sorting algorithm obtained from GA was assessed using real data. Results were compared with the sorting independently and blindly by three expert operators.

## 3. Results and discussion

### 3.1. The optimal values of t-SNE and DBSCAN parameters

According to GA results, the optimal values of t-SNE and DBSCAN parameters were obtained (Table 3). Based on these optimal values, the optimal spike sorting algorithm was developed and assessed for simulated and real datasets.

**Table 3.** the optimal values of t-SNE and DBSCAN parameters.

| Parameters: | Optimal values |
| --- | --- |
| t-SNE: |  |
| Distance Metric | cityblock |
| Perplexity | 70 |
| Exaggeration | 4 |
| Number of Dimension | 3 |
| DBSCAN: |  |
| $\epsilon$ | 3.39 |
| MinPts | 2.58 |

### 3.2. Spike sorting results

The results of the proposed algorithm for a sample population of 10 neurons are shown in Fig. 3. The optimal t-SNE and DBSCAN algorithm correctly detected 10 neurons, with accuracies ranging from 97.1% to 100.0% across neurons (98.6±0.1%). In this example, there were no false positive neurons. The sorting identification sensitivity and precision were on average 99.2±0.7 and 99.3±0.6, respectively. There was not any firing time inconsistency (i.e., ISI values less than 2 ms) in any of the detected classes. The overall performance of the robust algorithm is summarized in Table 4 and Figure 4. Overall, the number of missed and erroneously detected neurons was 1.5±1.3 and 1.4±0.5, respectively. The overall sensitivity, precision and accuracy for correctly identified neuron classes were 97.8±1.2, 90.1±4.8 and 94.4±1.4 respectively. These results indicate that the combination of t-SNE and DBSCAN works well for sorting spikes of different neurons in a fully automatic fashion. Next, the performance of our algorithm was compared with WAVECLUS using the same synthetic data.

**Figure 3.**
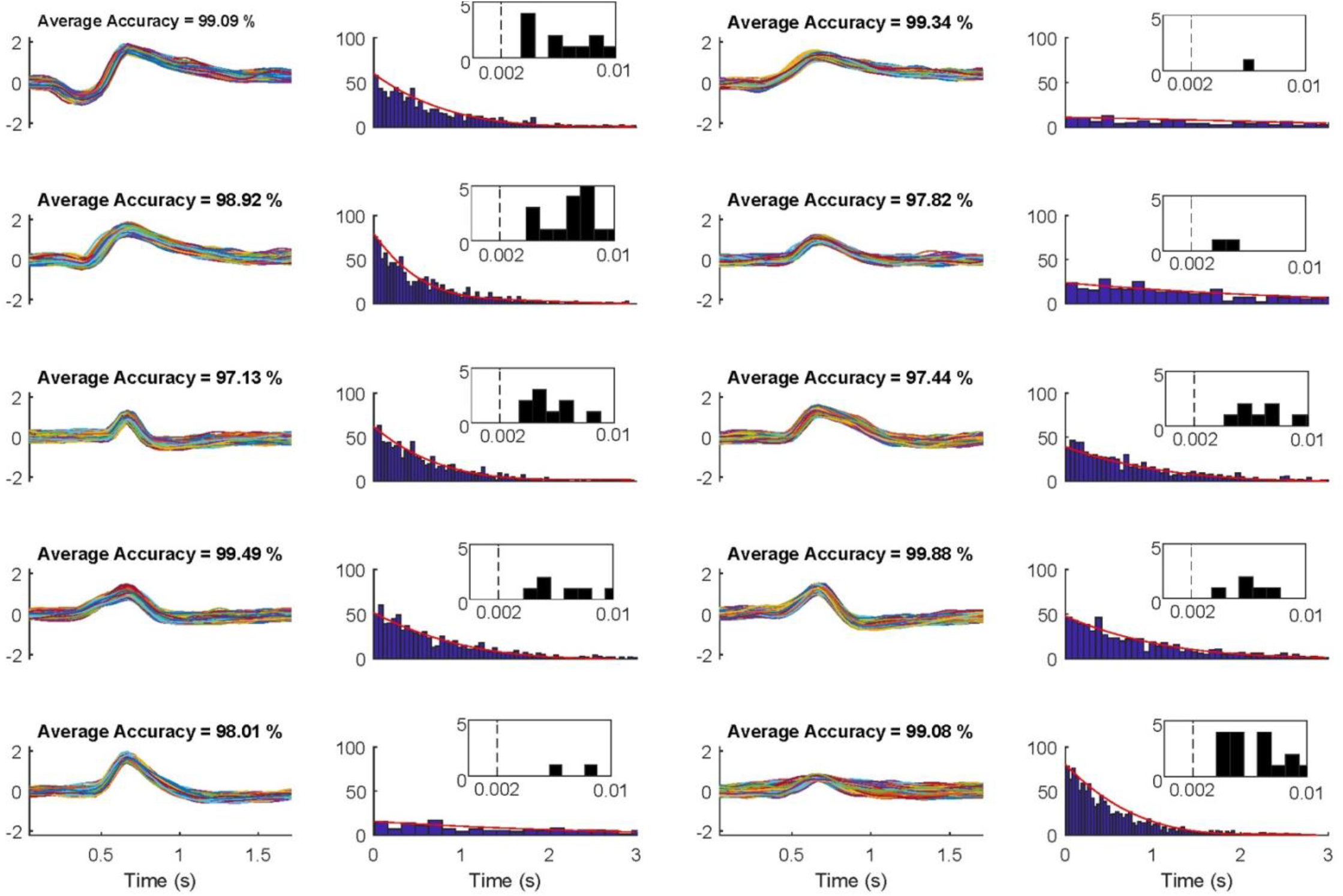
The spike shape and ISI of each neuron determined by our t-SNE and DBSCAN algorithm using GA optimized parameters on a session with 10 simultaneously neurons. There were 10 hit neurons with zero missed neurons. There were clusters and any refractory period violations. Performance for each neuron is shown in a pair of plots: all detected spike shapes are shown in the left plot and the ISI distribution is shown in the right plot. The zoomed in distribution of ISIs is shown as an inset in the right plot to allow one to examine ISIs less than 2ms refractory period.

**Table 4.** The performance of the proposed algorithm shown as mean±std.

| Number of<br>neurons | Clustering |  | Identification (%) |  |  |
| --- | --- | --- | --- | --- | --- |
|  | Hit | Miss | FP | TPR | PPV |
| 2 | 2.0±0.0 | 0.0±0.0 | 1.4±0.9 | 0.93±0.10 | 0.98±0.01 |
| 3 | 2.8±0.4 | 0.2±0.4 | 1.4±1.1 | 0.99±0.01 | 0.91±0.15 |
| 4 | 3.8±0.4 | 0.2±0.4 | 0.6±0.9 | 0.98±0.04 | 0.93±0.09 |
| 5 | 4.8±0.4 | 0.2±0.4 | 1.2±0.4 | 0.98±0.01 | 0.94±0.08 |
| 6 | 5.6±0.6 | 0.4±0.5 | 1.0±0.7 | 0.98±0.01 | 0.91±0.09 |
| 7 | 6.6±0.9 | 0.4±0.9 | 0.8±0.4 | 0.99±0.00 | 0.93±0.13 |
| 8 | 7.4±0.5 | 0.6±0.5 | 1.2±0.4 | 0.97±0.04 | 0.91±0.06 |
| 9 | 8.6±0.9 | 0.4±0.9 | 0.6±1.1 | 0.97±0.02 | 0.92±0.10 |
| 10 | 8.4±1.8 | 1.6±1.8 | 1.6±1.5 | 0.97±0.04 | 0.86±0.10 |
| 11 | 8.8±1.3 | 2.2±1.3 | 1.8±1.8 | 0.97±0.03 | 0.82±0.08 |
| 12 | 10.4±0.5 | 1.6±0.5 | 1.2±0.4 | 0.98±0.01 | 0.86±0.05 |
| 13 | 11.6±0.5 | 1.4±0.5 | 1.0±0.7 | 0.98±0.02 | 0.86±0.06 |
| 14 | 11.2±1.1 | 2.8±1.1 | 1.6±0.9 | 0.97±0.01 | 0.81±0.04 |
| 15 | 11.4±2.0 | 3.6±2.1 | 2.0±1.2 | 0.98±0.01 | 0.82±0.10 |
| 16 | 14.8±0.4 | 1.2±0.4 | 0.4±1.1 | 0.98±0.01 | 0.90±0.04 |
| 17 | 13.8±1.7 | 3.2±1.8 | 2.0±0.7 | 0.98±0.01 | 0.82±0.07 |
| 18 | 15.2±1.1 | 2.8±1.1 | 1.2±0.4 | 0.98±0.00 | 0.83±0.08 |
| 19 | 15.2±1.6 | 3.8±1.6 | 0.8±1.1 | 0.98±0.01 | 0.81±0.06 |
| 20 | 16.4±2.2 | 3.6±2.2 | 0.8±0.8 | 0.98±0.01 | 0.83±0.09 |

**Fig. 4.**
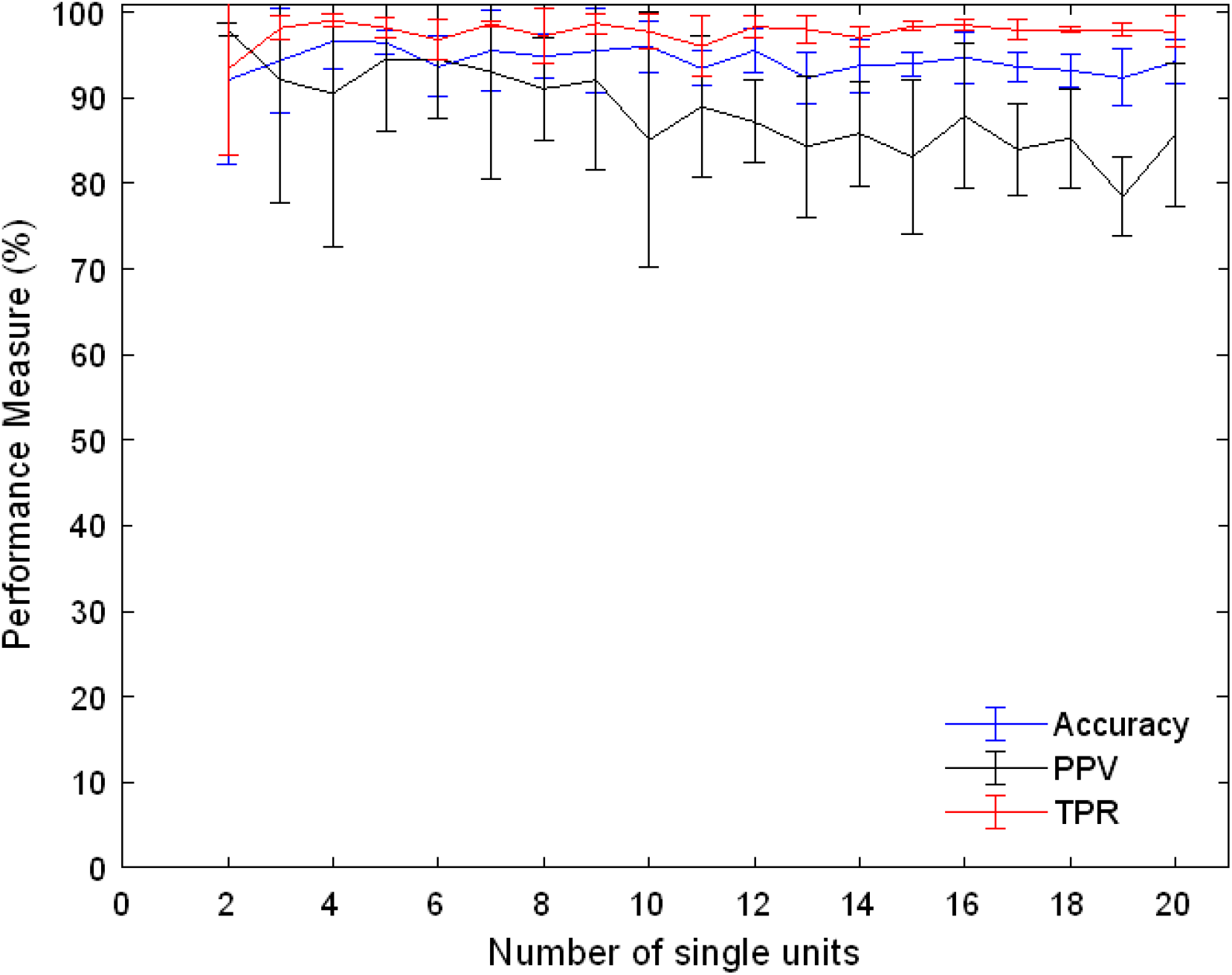
The accuracy, PPV and TPR performances of the proposed algorithm on all sessions with different number of simultaneously recorded neurons ranging from 2 to 20. The values shown are mean across sessions and the error-bars indicate std.

### 3.3. Comparison with the state-of-the-art

The sorting algorithm proposed by Quiroga et al. (2004), which is an open source and freely available software (https://github.com/csn-le/wave_clus), known as WAVECLUS, was used for comparison with the proposed optimal t-SNE and DBSCAN sorter algorithm. It has been shown in the literature that the other well-known algorithm, KlustaKwik, proposed by Harris et al. (2000), has the same performance as the WAVECLUS algorithm (Pedreira et al., 2012). They are, in fact, among the most cited algorithms in spike sorting literature (Wild et al., 2012). The neural decoded data used for comparison was the previously published result of the operation of three experts with parameter optimization using the WAVECLUS GUI (Pedreira et al., 2012).

The performance of the optimized t-SNE and DBSCAN sorter algorithm was compared with that of WAVECLUS in terms of the number of the units that were correctly identified (hits) as well as the maximum and minimum (Fig. 5). Note that we have taken the results of WAVECLUS as previously published in Quiroga et al. (2004) and have used the same performance measures for comparison. As seen in this figure, the average number of hits as well as maximum and minimum of the optimal t-SNE and DBSCAN sorter algorithm were overall higher than with those of WAVECLUS. There was a 48% increase in the number of hits using our algorithm compared with WAVECLUS. Although the two methods were almost similar when the number of neurons was less than or equal to 7, the proposed algorithm significantly outperformed the other algorithm when the number of neurons was greater than or equal to 8 (Fig. 5B).

**Figure 5.**
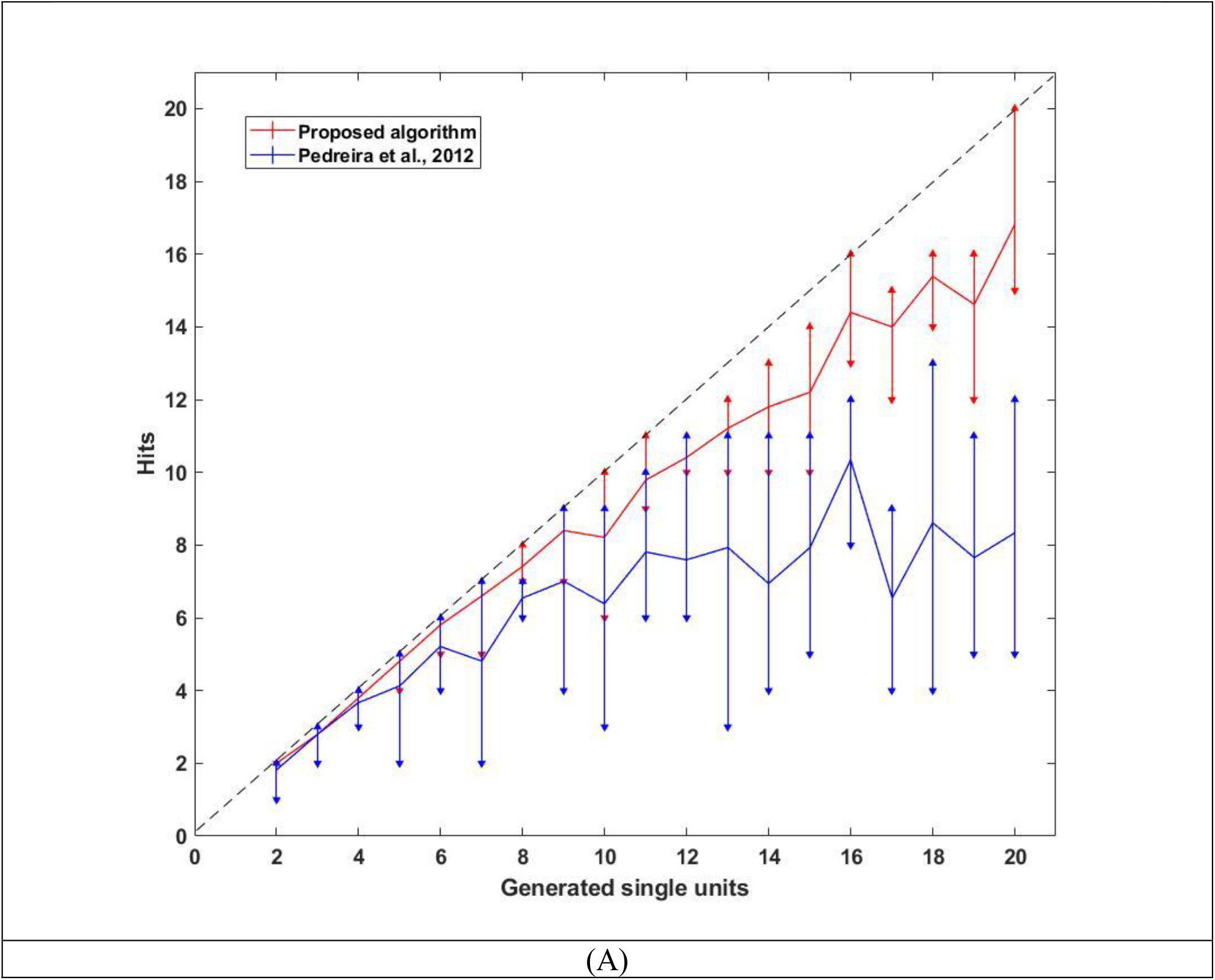
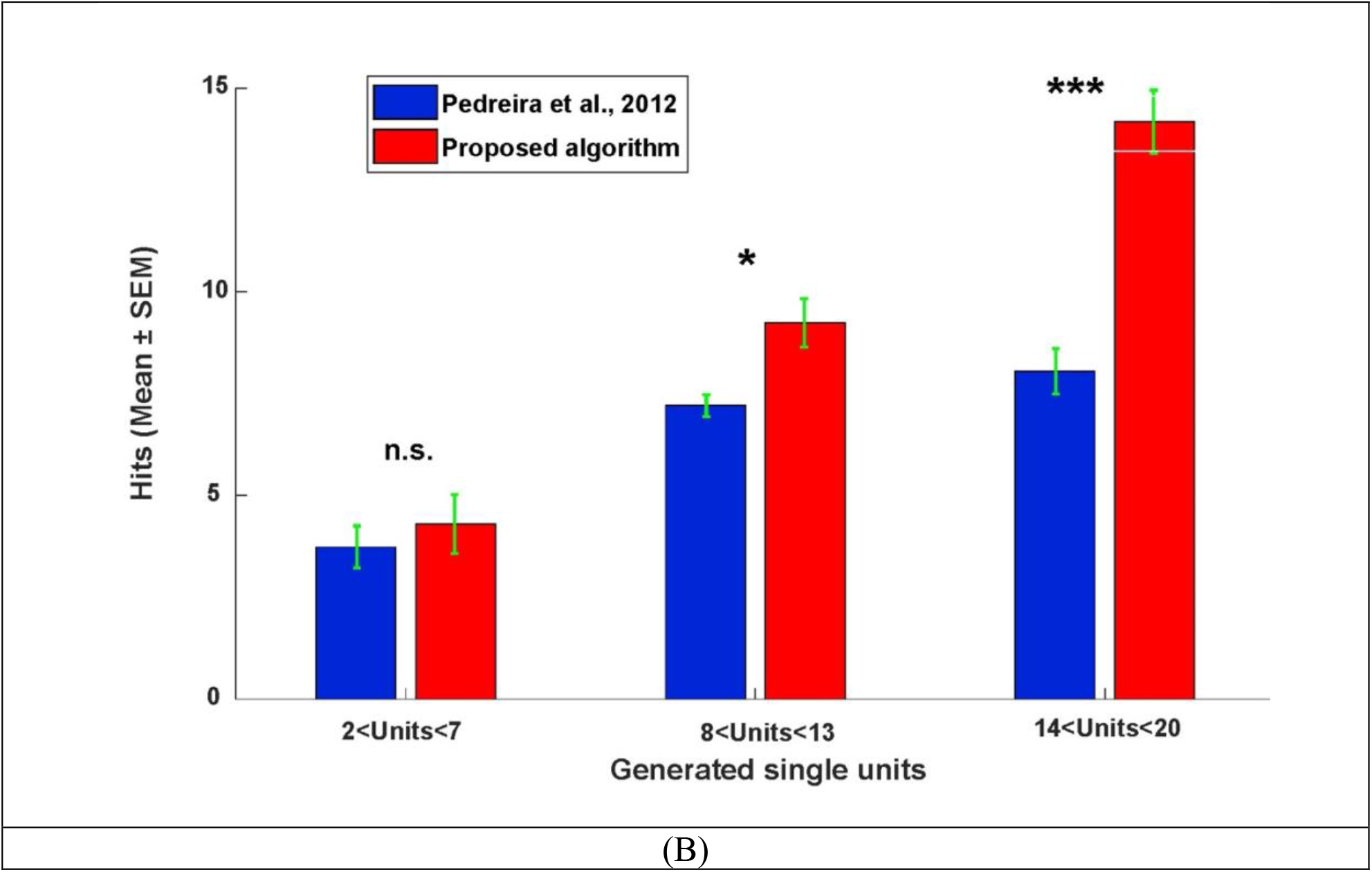
Comparison of performance of our model compared to WAVECLUS (Pedreira et al., 2012; Rey et al., 2015) on sessions from simulated data. (A) the number of hits with increasing number of neurons. Upward-pointing triangle and downward-pointing triangle denote maximum and minimum of hits, respectively. The dashed line has slope of one. (B) The average number hits for sessions with 2-7, 8-13 or 14-20 units. (mean ± SEM is shown, n.s., * and *** means nonsignificant, *p*-value< 0.05 and *p*-value< 0.001, respectively)

The average missed neuron and false positive errors of the proposed algorithm for different numbers of neurons is shown in Fig. 6. The proposed algorithm outperformed the WAVECLUS algorithm (Rey et al., 2015)(Fig. 5), particularly in terms of missed errors (see Fig. 4B of Pedreira et al. (2012)).

**Figure 6.**
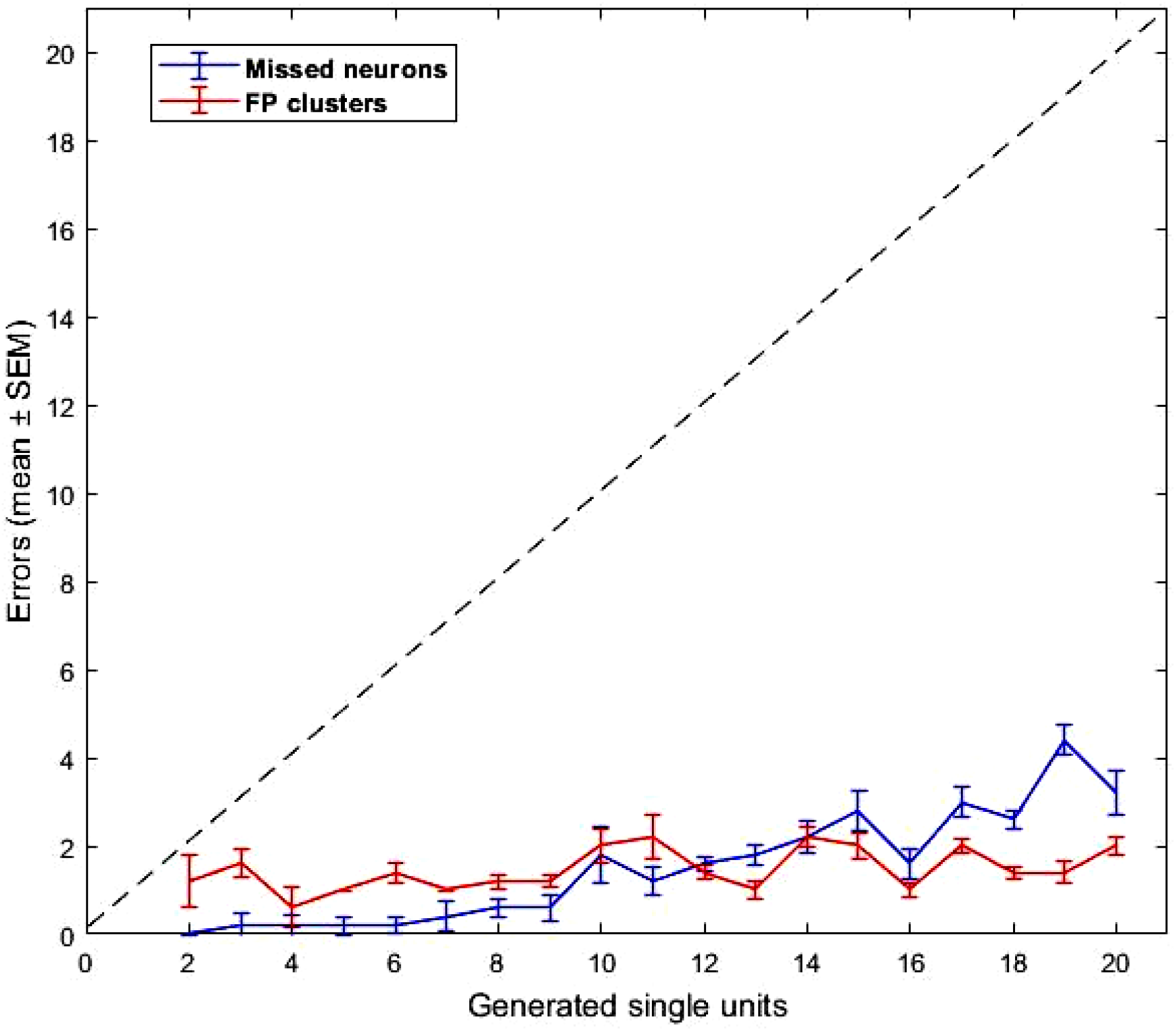
The Average number of missed neuron and false positives in the proposed algorithm. Error-bars denote SEM.

The knowledge derived from these results is that the proposed algorithm could effectively improve spike sorting of simulated data. To further validate our method, our algorithm was next tested on real data, with results presented in following section.

### 3.4. Comparison with real data

The performance of the optimal t-SNE and DBSCAN sorter algorithm on real data was compared with sorting carried out by three human experts using Plexon offline sorter V3.3.5 manually. For this purpose, 91691 spikes are detected using an amplitude threshold after filtering real data, and each expert clustered these spikes separately. The spikes were also separately clustered by our fully automated algorithm. Results of sorting the real data with the proposed algorithm are shown in Figure 7. Since there is no ground truth here for comparison and statistical analysis, we use the intersection of both spikes in corresponding clusters in the proposed algorithm and experts as a measure of truly detected spikes. Using this method, as seen in Fig. 8A, the corresponding spike clusters have an intersection area (*c*) of mutually detected spikes, as well as relative complements of spikes detected by the proposed algorithm, but not by experts (*a*), and vice versa (*b*).

**Figure 7.**
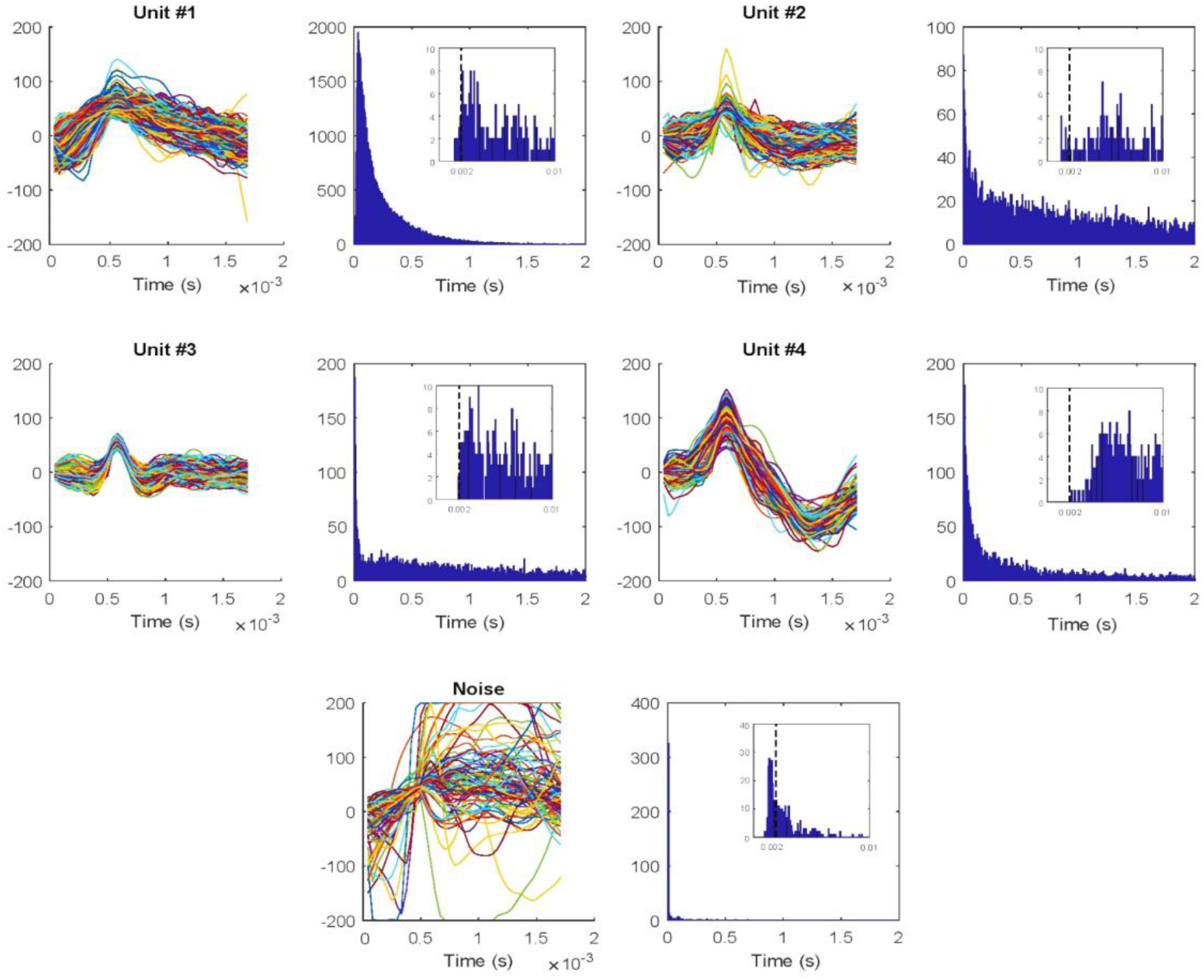
The spike shape and ISI of each neuron using our proposed optimal t-SNE and DBSCAN algorithm on a real signal with 4 neurons. Performance for each neuron is shown in a pair of plots: all detected spike shapes are shown in the left plot and the ISI distribution is shown in the right plot. The zoomed in distribution of ISIs is shown as an inset in the right plot to allow one to examine ISIs less than 2ms refractory period. Spike numbers with ISIs below the minimum refractory period (2 ms) is 0.02% (14 out of 68443 spikes) for unit #1, 0.03% (3 out of 8011 spikes) for unit #2, 0% (0 out of 6601 spikes) for unit #3 but is 32% (169 out of 509 shapes) for noise cluster.

As a measure of quality of spike sorting, we assumed that the average correlation between the spikes in relative complements in our algorithm and in the expert with the representative of spikes in the intersection area can tell us whether detected spikes in the relative complements represented true positives. The higher this average correlation, the better the sorting method (Table 5). We assume that this average correlation can speak to the homogeneity of detected spikes in each method, compared to their intersection. The homogeneity is defined as the average of correlation coefficients between representatives (mean point) of intersection cluster (c) with relative complement spikes in areas ‘a’ or ‘b’. The same measure was calculated by comparing spike clusters between experts. As shown, the homogeneity of responses in the proposed algorithm was not significantly different from that of experts (Fig 8B, F_2,33_=2.45 P=0.09), however there was a nonsignificant trend towards marginally higher homogeneity for the algorithm.

**Table 5.** The performance of the proposed algorithm vs. experts spikes sorting.

| Cluster | #1 | #2 | #3 | #4 | Noise |
| --- | --- | --- | --- | --- | --- |
| **Expert #1** |  |  |  |  |  |
| **Spikes numbers** |  |  |  |  |  |
| Proposed algorithm | 68443 | 8127 | 8011 | 6601 | 509 |
| Expert | 56096 | 2426 | 7639 | 6234 | 19296 |
| intersection of Expert and proposed algorithm | 56042 | 2328 | 7052 | 6233 | 472 |
| In expert-Not in proposed algorithm | 54 | 98 | 587 | 1 | 18824 |
| In proposed algorithm - Not in expert | 12401 | 5799 | 959 | 368 | 37 |
| **Average of $\rho$ between intersection clusters with** |  |  |  |  |  |
| In expert-Not in proposed algorithm | 85.74 | 78.86 | 75.03 | 74.46 | - |
| In proposed algorithm - Not in expert | 86.19 | 87.87 | 86.59 | 97.77 | - |
| **Percentage of spikes numbers with $\rho < 90\%$** |  |  |  |  |  |
| In expert-Not in proposed algorithm | 64.81 | 92.86 | 96.42 | 100 | - |
| In proposed algorithm - Not in expert | 59.1 | 56.91 | 72.89 | 2.18 | - |
| **Expert #2** |  |  |  |  |  |
| **Spikes numbers** |  |  |  |  |  |
| Proposed algorithm | 68443 | 8127 | 8011 | 6601 | 509 |
| Expert | 63211 | 5181 | 9846 | 5700 | 7753 |
| intersection of Expert and proposed algorithm | 63153 | 3728 | 7345 | 5696 | 471 |
| In expert-Not in proposed algorithm | 58 | 1453 | 2501 | 4 | 7284 |
| In proposed algorithm - Not in expert | 5290 | 4399 | 666 | 905 | 38 |
| **Average of $\rho$ between intersection clusters with** |  |  |  |  |  |
| In expert-Not in proposed algorithm | 88.01 | 88.12 | 72.94 | 80.01 | - |
| In proposed algorithm - Not in expert | 84.5 | 86.61 | 87.09 | 99.34 | - |
| **Percentage of spikes numbers with $\rho < 90\%$** |  |  |  |  |  |
| In expert-Not in proposed algorithm | 63.79 | 81.21 | 96.8 | 50 | - |
| In proposed algorithm - Not in expert | 75.88 | 61.53 | 83.63 | 0.11 | - |
| **Expert #3** |  |  |  |  |  |
| **Spikes numbers** |  |  |  |  |  |
| Proposed algorithm | 68443 | 8127 | 8011 | 6601 | 509 |
| Expert | 55199 | 5344 | 7012 | 5926 | 18210 |
| intersection of Expert and proposed algorithm | 55176 | 3956 | 7001 | 5895 | 406 |
| In expert-Not in proposed algorithm | 23 | 1388 | 11 | 31 | 17804 |
| In proposed algorithm - Not in expert | 13267 | 4171 | 1010 | 706 | 103 |
| **Average of $\rho$ between intersection clusters with** |  |  |  |  |  |
| In expert-Not in proposed algorithm | 85.63 | 86.46 | 80.12 | 82.00 |  |
| In proposed algorithm - Not in expert | 92.50 | 86.72 | 90.51 | 98.72 |  |
| **Percentage of spikes numbers with $\rho < 90\%$** |  |  |  |  |  |
| In expert-Not in proposed algorithm | 52.18 | 70.24 | 72.73 | 64.52 |  |
| In proposed algorithm - Not in expert | 22.74 | 31.41 | 43.66 | 0.57 |  |

As another quality measure, we defined consistency as the number of spikes in the intersection clusters (c) divided by spike numbers of ‘a’∪̛c’ or spike numbers of ‘c’∪̛b’ expressed as a percentage (Fig 8C). As shown in Fig. 8C, the consistency between experts and the proposed method is significantly higher than the consistency among experts themselves, meaning that the algorithm is consistent with the consensus of the experts.

**Figure 8.**
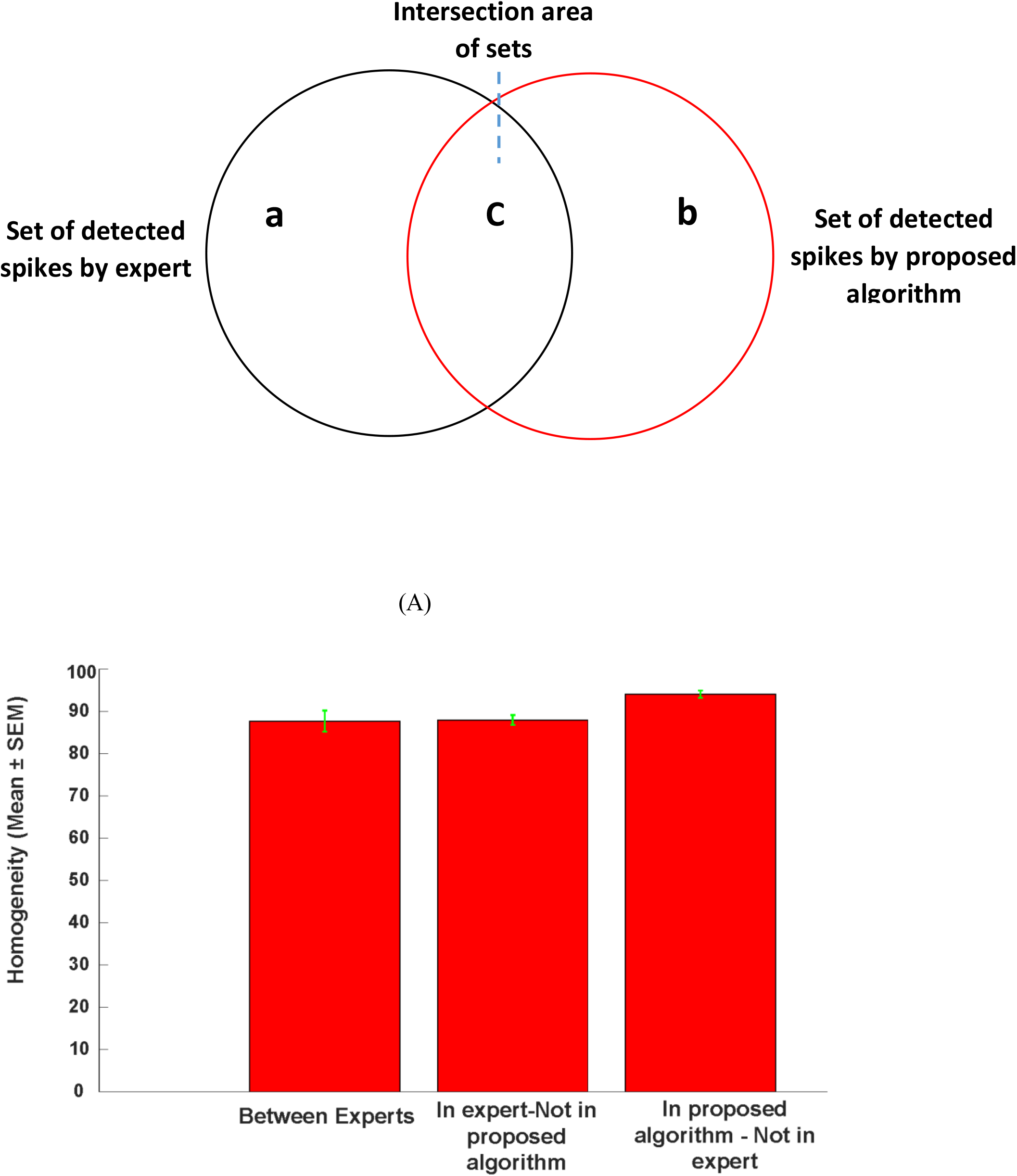
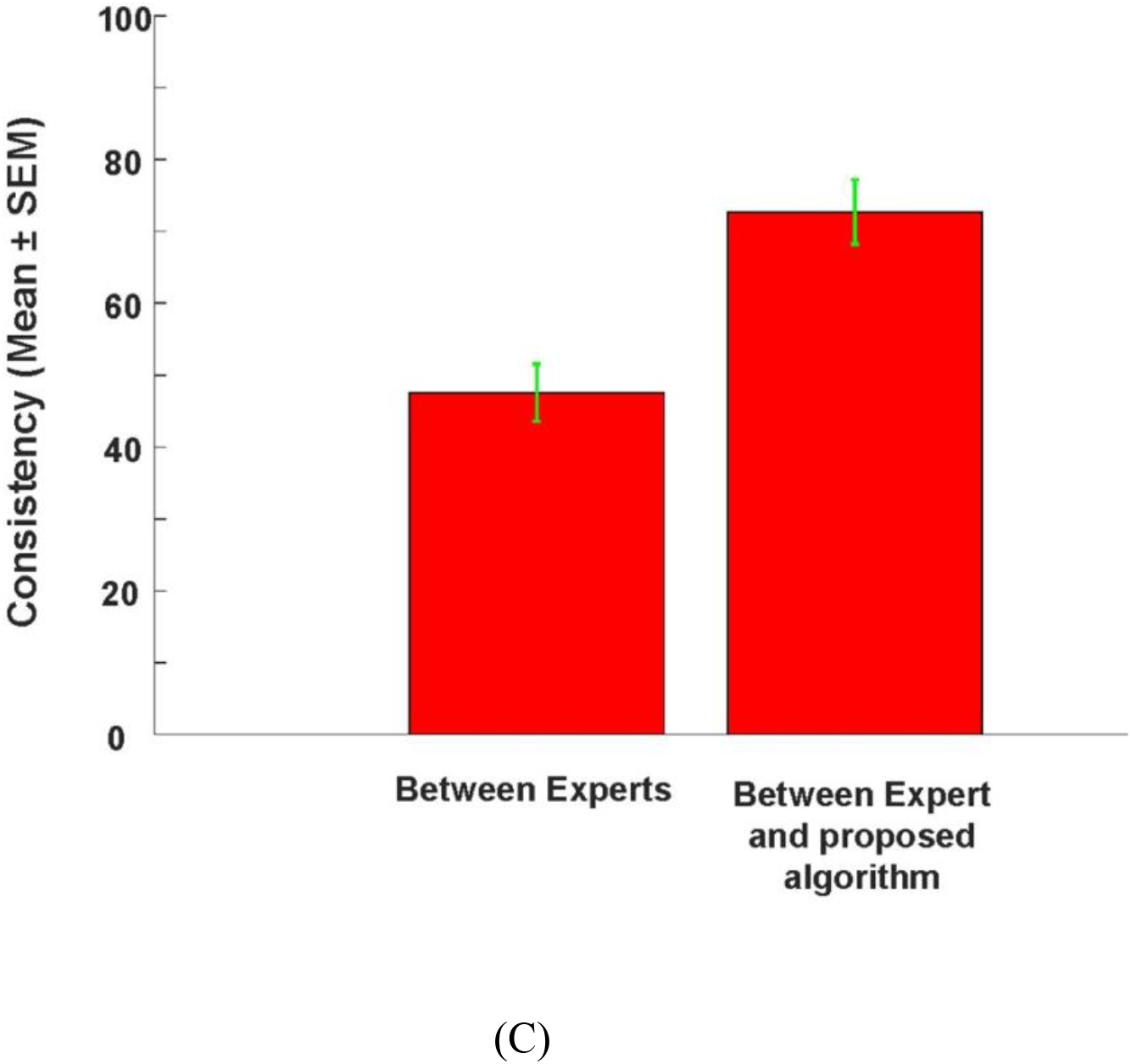
Comparison spike sorting performance of our algorithm with that of 3 experts on real data (A) The Venn diagram showing the relationship between detected spikes for a given unit using our algorithm and a given expert area ‘c’ (intersection area) denotes spikes detected by both methods. Areas ‘a’ show spikes detected by the expert not our algorithm and areas ‘d’ denote spikes detected by our algorithm not by the expert (complement areas). (B) Average of Pearson’s correlation coefficient in the complement areas between experts, and for the complement of in expert – not in proposed algorithm and for the complement of proposed algorithm – not in expert. The higher this average correlation coefficient in the complement areas indicated the higher the homogeneity of detected spikes (C) (C) Average percentage of detected spikes in the intersection area ‘c’ compared with all detected spikes when comparing between experts and between experts and our algorithm. This percentage shows the consistency of various spike detection methods. Consistency of our algorithms and experts was higher than consistency between experts
**Homogeneity:** Average of correlation coefficients between representative of intersection clusters (*C*) with relative complements spikes A ∪ B or C ∪ D
**Consistency:** Spikes numbers of intersection clusters (*C*) divides by Spikes numbers of A ∪ B ∪ C or Spikes numbers of C ∪ D ∪ E (based on percentage)

According to Table 5, the average Pearson correlation coefficient between the representative of intersection cluster and relative complements of proposed algorithm (area ‘b’) #1-3 were 89.6, 89.4 and 92.11, respectively. These values for relative complements of experts (area ‘a’) were 78.52, 82.27 and 83.55, respectively. Thus, on average the correlation coefficient with intersection (area ‘c’) was higher for the algorithm compared to the experts.

Also, we assumed that percentage of spikes with *ρ* < 90%, can be interpreted as false positive error. This percentage was also lower for the proposed algorithm compared to the experts. As seen in Table 5, the average of these values for experts #1-3 are 88.52, 72.95 and 64.91 versus 47.77, 55.28 and 24.59 for the proposed algorithm, respectively. As a result, it is clear that the combination of t_SNE with DBSCAN algorithm considerably improved the sorting of real data.

## 4. Discussion

Direct electrophysiological recording of neurons is the gold standard for understanding signal processing in the brain. Recent advances in technology that allow simultaneous recording of many neurons across a large number of channels simultaneously present a challenge for manual spike sorting. In this study, a combination of two methods, t-SNE and DBSCAN, was developed for an off-line and fully-automated spike sorting algorithm. Our algorithm outperformed the state of the art methods for spike sorting using WAVECLUS, as well as manual spike sorting by experts in simulated and real neural recordings. Specifically, our algorithm performance was significantly better than WAVECLUS when the number of neurons was large (Fig 5); the sensitivity and accuracy of spike sorting was above 90% and specificity was above 80% in simulated data for up-to 20 simultaneously recoded neurons (Fig 4). Detected neurons had distinct spike shapes with ISI distribution outside the refractory period in almost all cases in both simulated (Fig 3) and real data (Fig 7). Comparison of algorithm performance with that of manual sorting by experts showed equal or better performance as measured by homogeneity of spike shapes for detected neurons (Fig 8b).

The six parameters in our algorithm were optimized using a genetic algorithm. While this algorithm was optimized on the simulated data, using the same parameters on the real spike seemed to give satisfactory results compared to the manual sorting by experts. Ideally, if manual sorting for a large number of different experiments become available the parameter optimization can be done on matching performance of the experts while maximizing the desired cost function such as homogeneity or consistency. Such optimization should result in even better performance of the algorithm on real data in the future.

One limitation of spike sorting that we tried to address here was the sorting of big numbers of neurons. Although some methods tried to tackle this issue (Ekanadham et al., 2014; Pedreira et al., 2012), most of the sorting methods to date have focused on sorting small numbers of neurons (< 10) (Carlson et al., 2014; Franke et al., 2010). One of the advantages of our method is its ability to deal with sparse firing neurons while most of the other algorithms (Ekanadham et al., 2014; Franke et al., 2010; Hilgen et al., 2017; Yger et al., 2016) miss these neurons because of few spikes per second. This is because in those algorithms neurons with low firing rate are often discarded as noise or a grouped together with neurons with more numerous but similar spikes. A problem that is avoided in our algorithm by using density based clustering which is less sensitive to the actual cluster shape and size. The other advantage of the proposed algorithm is that it is fully automated without the need for manual post processing correction. As sorting of large signals or big number of channels can be difficult and cumbersome, having an automatic and unsupervised method compared to other supervised or semi-supervised algorithms (Adamos et al., 2008; Calabrese and Paninski, 2011; Haga et al., 2013; Kim and Kim, 2003; Quiroga et al., 2004; Vargas-Irwin and Donoghue, 2007) would be highly desirable.

Results highlight advantages of our proposed algorithm in sorting data from brain regions as well as a simulated dataset using the same parameters, which illustrates the power of this approach. Results also indicated that the t-SNE method handles some changes in waveforms of spikes, which may result from movement of electrodes relative to the tissue. We have provided a software equipped with a graphical user interface (GUI) that implements our t-SNE-DBSCAN algorithm along with this paper. Although our software runs quickly on datasets with low numbers of spikes, the clustering time theoretically scales linearly with the number of spikes.

Taken together, our results demonstrate the optimal t-SNE and DBSCAN sorter algorithm can perform spike sorting in a fully automated fashion with high accuracy, sensitivity and precision. In future work, we will be extending our algorithm to handle cases such as tetrode recording in which the same unit can appear in multiple channels. In addition, the current algorithm will take some time to sort the recorded spikes into separate clusters corresponding to each unit and thus is not well suited for online applications. Adaptations of this model where clustering can be done adaptively and in a trial-by-trial fashion in real time would be an important extension for future work, which will allow its use in brain machine interface applications as well.

## Acknowledgment

The authors acknowledge and appreciate the funding support provided by the IPM. The authors would like to thank Dr. Bahareh Taghizadeh for helpful comments.

